# Gamma band alterations and REM-like traits underpin the acute effect of the atypical psychedelic ibogaine

**DOI:** 10.1101/2020.06.25.172304

**Authors:** Joaquín González, Matias Cavelli, Santiago Castro-Zaballa, Alejandra Mondino, Adriano BL Tort, Nicolás Rubido, Ignacio Carrera, Pablo Torterolo

## Abstract

Ibogaine is a psychedelic alkaloid that has attracted scientific interest because of its important antiaddictive properties evidenced in observational studies in humans, and in models for substance-use-disorders in rodents. Its subjective effect has been described as intense vivid dream-like experiences occurring while awake; hence, ibogaine is often referred to as an oneirogenic psychedelic. While this unique dream-like profile has been hypothesized to aid the antiaddictive effects in the past, the electrophysiological signatures of the ibogaine psychedelic state remain unknown. In our previous work, we showed in rats that ibogaine administration promotes a waking state with abnormal motor behavior, accompanied by a decrease in NREM and REM sleep. Here, we performed an in-depth analysis of the intracranial electroencephalogram during “ibogaine wakefulness”. Ibogaine induced gamma oscillations with larger power than control levels but less coherent and less complex; i.e., this state shows clear REM sleep traits within the gamma frequency band. Thus, our results provide novel biological evidence for the association between the psychedelic state and REM sleep, and an empirical basis for the oneirogenic conjecture of ibogaine.

## INTRODUCTION

Ibogaine is a potent psychedelic alkaloid that has attracted scientific interest because of its long-lasting antiaddictive properties, evidenced in anecdotal and observational studies in humans^1–5^, and in an extensive pre-clinical work in rodents^6,7^. Subjective reports portray the ibogaine experience as entering into an intense dream-like episode while awake, involving memory retrieval and prospective imagination, without producing the typical interferences in thinking, identity distortions, and space–time alterations produced by classical psychedelics (e.g. DMT, LSD, psilocybin)^8–14^. Thus, ibogaine is often referred as an oneirogenic psychedelic^8,10,15^.

In spite of the vast amount of research regarding the antiaddictive effects of ibogaine, the biological substrate of its unique oneiric effects remains elusive. Although seemingly unrelated, the oneirogenic effects of ibogaine have been hypothesized to aid its antiaddictive properties^6,16^. Taking into account that most vivid dreams occurs during REM sleep, the dream-like experiences would be the manifestation of a REM sleep-like brain state, which in turn could favor the antiaddictive effects through an increase in neural plasticity and memory reconsolidation, similar to previously reported functions of natural REM sleep^17^. Therefore, if this conjecture is true, we should expect to find REM sleep characteristics in the electrocortical activity following the administration of ibogaine.

In our previous work^18^, we showed in rats that ibogaine promotes a wakefulness state with abnormal motor behaviors in a dose dependent manner. These effects were accompanied by a decrease in NREM sleep and a profound REM sleep suppression. Nevertheless, as the analysis relied on visual inspection, we were not able to answer which features characterize the waking state induced by ibogaine. Therefore, in the present work we employed a state-of-the-art computational analysis of the intracranial electroencephalogram (iEEG) to analyze the acute effects of ibogaine. We found a unique iEEG profile, which differs from that of physiological wakefulness and is compatible with a REM-like brain state. Hence, our results provide the first electrophysiological evidence of a dream-like brain state produced by ibogaine.

## MATERIAL AND METHODS

### Ibogaine

Ibogaine was obtained and purified from *T. Iboga* extracts following the procedures employed in ^18^. A 40 mg/kg dose (i.p.) was employed in this work. This dose has the largest effect on sleep and motor behavior^18^ and correlates with the high doses involved in shamanic and medicinal uses of ibogaine^19^.

### Experimental Animals

Six Wistar adult rats were maintained on a 12-h light/dark cycle (lights on at 07.00 h). Food and water were freely available. The animals were determined to be in good health by veterinarians of the institution. All experimental procedures were conducted in agreement with the National Animal Care Law (No. 18611) and with the “Guide to the care and use of laboratory animals” (8th edition, National Academy Press, Washington DC, 2010). Furthermore, the Institutional Animal Care Committee approved the experimental procedures (Exp. N° 070153-000332-16). Adequate measures were taken to minimize pain, discomfort, or stress of the animals, and all efforts were made to use the minimal number of animals necessary to obtain reliable scientific data.

### Surgical Procedures

The animals were chronically implanted with electrodes to monitor the states of sleep and wakefulness. We employed similar surgical procedures as in our previous studies^18,20,21^. Anesthesia was induced with a mixture of ketamine-xylazine (90 mg/kg; 5 mg/kg i.p., respectively). The rat was positioned in a stereotaxic frame and the skull was exposed. To record the iEEG, stainless steel screw electrodes were placed in the skull above motor, somatosensory, visual cortices (bilateral), the right olfactory bulb, and cerebellum, which was the reference electrode (see Table S1). To record the electromyogram (EMG), two electrodes were inserted into the neck muscle. The electrodes were soldered into a 12-pin socket and fixed onto the skull with acrylic cement. At the end of the surgical procedures, an analgesic (Ketoprofen, 1 mg/kg, s.c.) was administered. After the animals had recovered from these surgical procedures, they were left to adapt in the recording chamber for 1 week.

### Experimental Sessions and Sleep Scoring

Animals were housed individually in transparent cages (40 x 30 x 20 cm) containing wood shaving material in a temperature-controlled (21–24 degrees Celsius) room, with water and food *ad libitum*. Experimental sessions were conducted during the light period, between 10 AM and 4 PM in a sound-attenuated chamber with Faraday shield. The recordings were performed through a rotating connector, to allow the rats to move freely within the recording box. Polysomnographic data were acquired and stored in a computer using the Dasy Lab Software employing 1024 Hz as a sampling frequency and a 16 bits AC converter.

The states of sleep and wakefulness were determined in 10-s epochs. Wakefulness was defined as low-voltage fast waves in the motor cortex, a noticeable theta rhythm (4–7 Hz) in the somatosensory and visual cortices, and relatively high EMG activity. NREM sleep was determined by the presence of high-voltage slow cortical waves together with sleep spindles in frontal, parietal, and occipital cortices associated with a reduced EMG amplitude; REM sleep as low-voltage fast frontal waves, a regular theta rhythm in the occipital cortex, and a silent EMG except for occasional twitches. Artifacts and transitional epochs were removed employing visual supervision.

### Data Analysis

To evaluate the ibogaine effect on iEEG activity, we selected the first two hours following the ibogaine i.p. administration since almost continuous wakefulness and abnormal motor and autonomic effects (tremor, piloerection) were only evident during this period^18^. From the first two hours, only artifact-free wake epochs were analyzed from both the control and ibogaine experiments. NREM sleep epochs were selected from the entire 6 hours due to the reduced time of this state after ibogaine i.p. administration. Additionally, REM sleep epochs from control experiments were also examined. REM sleep following ibogaine administration was not considered due to the lack of this state in several animals.

### Power spectrum

The power spectrum was obtained by means of the *pwelch* built-in function in Matlab (parameters: window = 1024, noverlap = [], fs = 1024, nfft = 1024), which corresponds to 1-s sliding windows with half-window overlap, and a frequency resolution of 1 Hz. The time-frequency spectrograms were obtained employing the function *mtspecgramc* from the Chronux toolbox^23^ (available at: http://chronux.org), using 5 tapers and a time-bandwidth product of 5. All spectra were whitened by multiplying the power at each frequency by the frequency itself, thus counteracting the 1/f trend. In addition, the spectra were normalized to obtain the relative power by dividing the power value of each frequency by the sum across frequencies. The traditional frequency bands depicted in the figures were taken as: delta (1-4 Hz), theta (5-10 Hz), sigma (11-14 Hz), beta (15-29 Hz) and gamma (30-100 Hz).

### Spectral coherence

To measure synchronization between electrodes, we employed the magnitude squared coherence using the *mscohere* built-in function in Matlab (parameters: window = 1024, noverlap = [], fs = 1024, nfft = 1024), which corresponds to 1-s sliding windows with half-window overlap, and a frequency resolution of 1 Hz. The time-frequency coherograms were obtained employing the function *cohgramc* from the Chronux toolbox, using 10 tapers and a time-bandwidth product of 100.

### Cluster-based permutation test

To obtain statistical thresholds for group comparisons of power and coherence, we employed a data-driven approach comparing empirical clusters of frequencies instead of comparing traditionally defined frequency bands. The method consisted of first comparing individual frequencies (512 frequencies) in each condition by means of paired t-tests (alpha = 0.05). Once we obtained the p values for each frequency, all consecutive significant frequencies were grouped into empirical clusters (defining a minimum cluster size of 4 frequency points), and a new statistic was formed by summing the t-statistic of each frequency inside the cluster. To assess whether a given cluster was significant, a null hypothesis distribution of cluster statistics was constructed by randomizing labels (control and ibogaine) and repeating the cluster construction method for a total of 10000 randomizations. The p values of the empirical clusters were obtained by comparing each cluster statistic to the randomized cluster statistic distribution (X). We employed two-tailed comparisons for the power spectrum and permutation entropy (pvalue = 2min(P(X > Xobs)),P(X < Xobs)), and one-tailed for the coherence comparisons (pvalue = P(X < Xobs)).

### Permutation entropy

Prior to quantifying the permutation entropy, the iEEGs were down-sampled to 128 Hz. The framework consisted of 2 main steps. In the first step, we encoded the time-series into ordinal patterns (OP) following Bandt and Pompe method^24^. The encoding involves dividing a time-series {*x*(*t*), *t*=1,…,*T*} into ⌊(*T*−*D*)/*D*⌋ non-overlapping vectors, where ⌊y⌋ denotes the largest integer less than or equal to y and D is the vector length, which is much shorter than the time-series length (D≪T). Then, each vector is classified according to the relative magnitude of its D elements. Namely, we determined how many permutations between neighbors are needed to sort its elements in increasing order; then, an OP represents the vector permutations. The second step consists in applying the Shannon entropy to quantify the average randomness (information content) of the OP distribution. Shannon entropy is defined as H =−∑p(OP) log[p(OP)], where p(OP) is the probability of finding a given OP in the signal (among the set of all OPs), and the summation is carried over all possible OPs. To assess the statistical significance between conditions, we employed paired two-tailed t-tests with α =0.05.

### Sleep scoring neural network

A multi-layer perceptron (10 hidden layers) was employed to distinguish between the states of wakefulness and REM sleep. We used the built-in classification network *patternnet* in Matlab. The input to the network consisted of values of gamma power (OB, M1r, M1l, S1r) and coherence (the 9 significant pairs in Figure 2C). The network was trained through a supervised scheme employing the visually scored states in the control condition (either Wake or REM). The training was performed employing the scaled conjugate gradient backpropagation algorithm (*trainscg* built-in function in Matlab), and the performance of the network was evaluated by the cross-entropy algorithm (*crossentropy* built-in function in Matlab).

**Figure 1.**
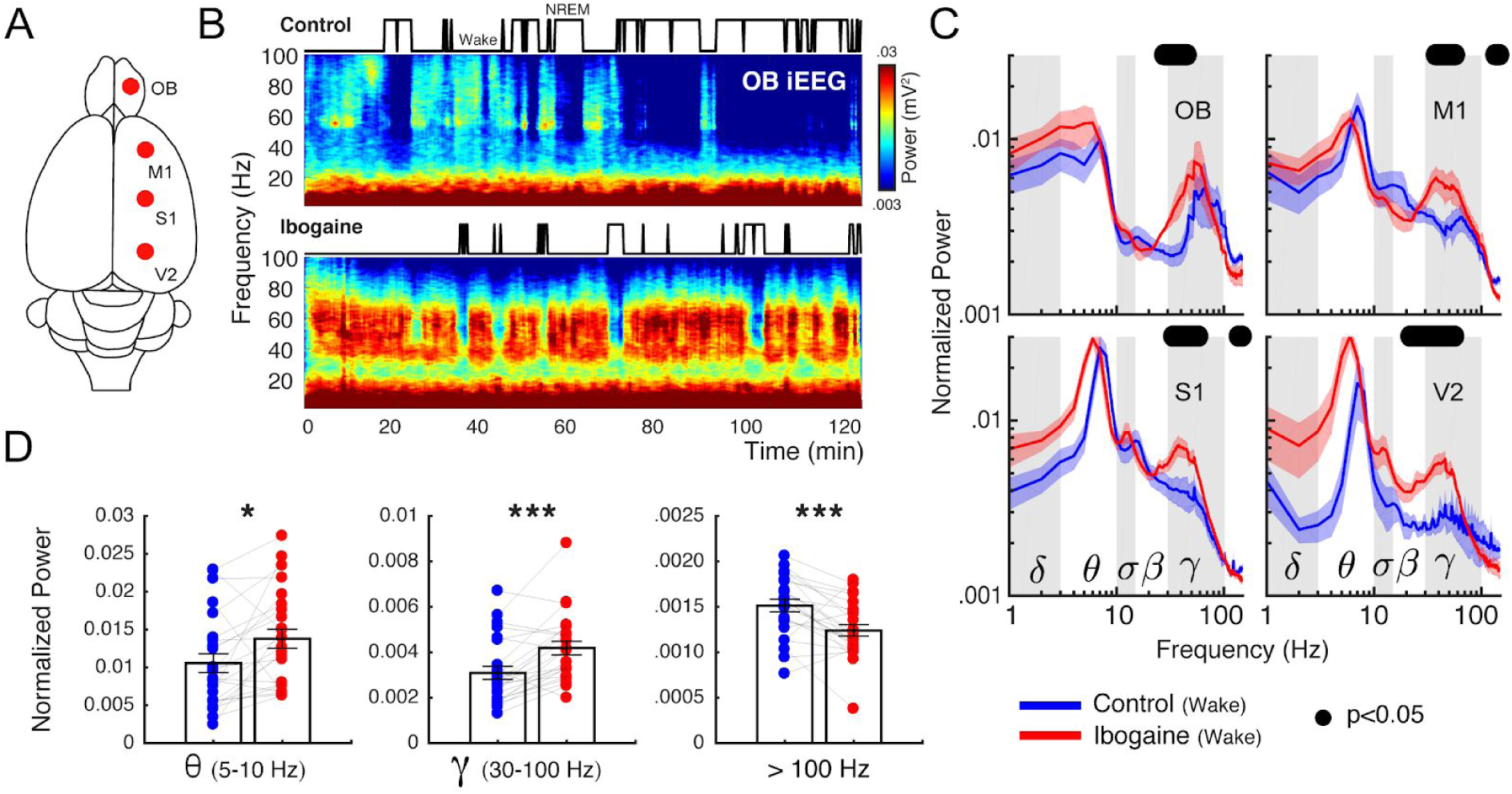
Ibogaine significantly alters iEEG frequency distribution. **A** Location of the analyzed intracranial electrodes in the right hemisphere (OB, olfactory bulb; M1, primary motor cortex; S1, primary somatosensory cortex; V2, secondary visual cortex). **B** Spectrograms from a representative animal following the administration of saline (control) and ibogaine (40 mg/kg). The hypnograms are plotted on top. **C** Normalized power spectra during wakefulness (see Material and Methods for normalization details). The solid line represents the mean (n=6 animals) for the first 2 hours post-injection; the shaded area depicts the standard error of the mean (S.E.M.). The black dots mark the statistically significant frequencies (p<0.05) corrected by a cluster-based permutation test. The traditional frequency bands (Greek letters) are delimited by gray and white boxes in each plot. The differences between hemispheres were minimal (see Figure S2). **D** Mean power for theta, gamma, > 100 Hz (up to 512 Hz) frequency bands. Each point corresponds to an electrode of a single animal; bars show mean ± S.E.M.. *p<0.05,*** p<0.001, paired t-test.

**Figure 2.**
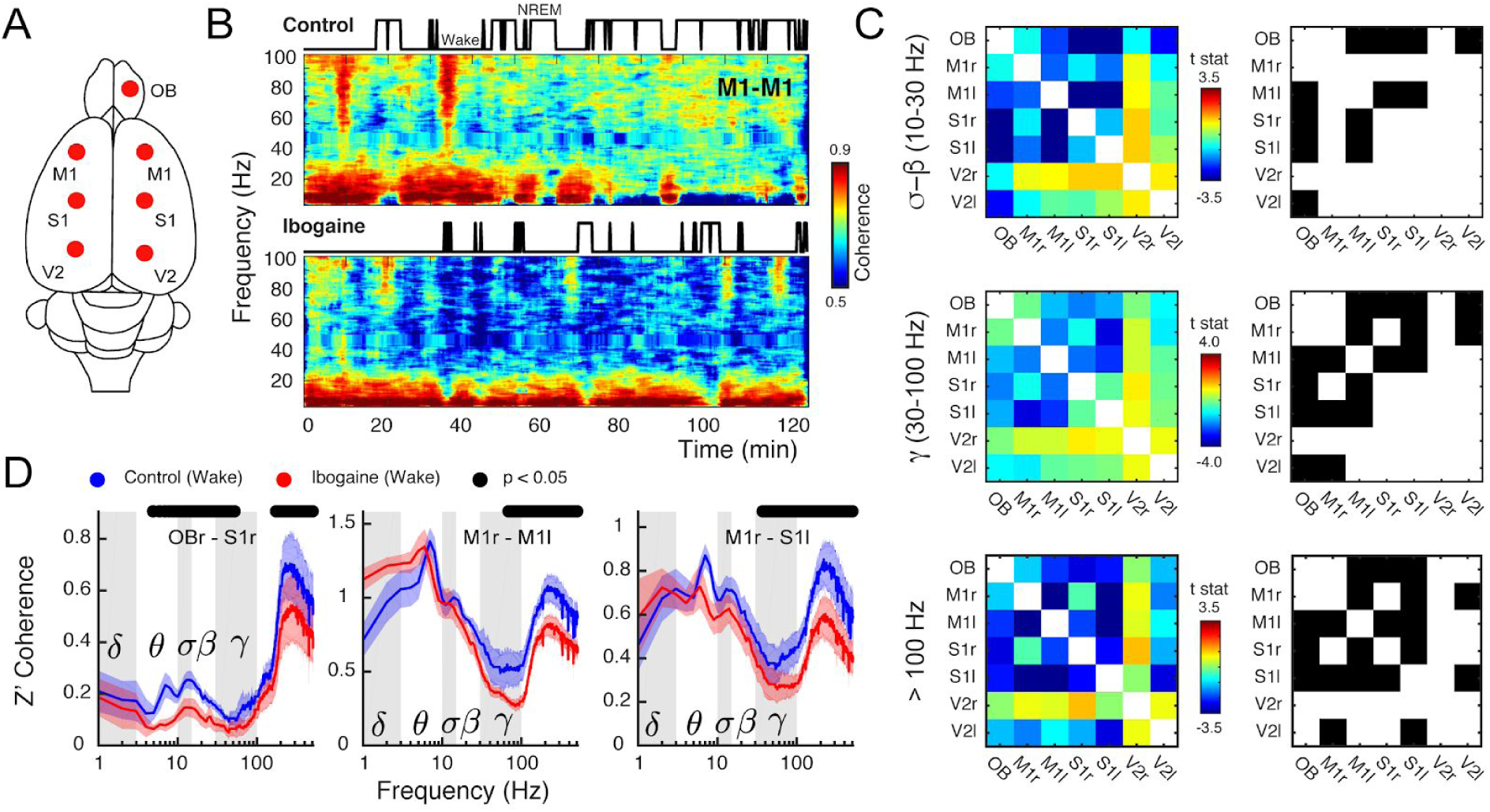
Ibogaine decreases long-range phase synchronization. **A** Location of the analyzed intracranial electrodes. **B** Coherogram following saline (control) and ibogaine (same animal and epoch as in Figure 1B). The hypnograms are plotted on top. This plot shows the phase coherence between the right and left primary motor cortex as a function of time and frequency. **C** The left column shows the t-statistic (t-stat) of the pair-wise coherence difference matrix (i.e., the average difference is divided by the S.E.M.) for three frequency bands (sigma-beta, gamma and > 100 Hz, up to 512 Hz). The right column shows the electrode pairs with a significant difference (p<0.05, corrected cluster-based permutation test; r: right; l: left). **D** Z’ coherence as a function of frequency of three representative combination of electrodes (same labels, statistical analysis, and wakefulness epochs as in Figure 1C).

### Phase-amplitude coupling

To measure coupling between frequencies within a same region, we employed the modulation index method^25^. Briefly, the raw signal was filtered between 1 and 15 Hz in 1 Hz steps (*eegfilt* function EEGLAB^26^; bandwidth 3 Hz) to obtain the slow frequency components, and then the phase time series were extracted from their analytical representation based on the Hilbert transform (*hilbert* bulit-in function in Matlab). In addition, the same raw signal was also filtered between 40 and 180 Hz in 10 Hz steps (bandwidth 10 Hz) to obtain the faster frequency components, and their amplitude time series are also obtained from the analytical representation. Then, phase-amplitude distributions were computed between all slow-fast frequency combinations. Finally, the modulation index was obtained as MI = (Hmax-H)/Hmax, where Hmax is the maximum possible Shannon entropy for a given distribution (log(number of bins)) and H is the actual entropy. The MI value of each slow-fast frequency combination was plotted in pseudocolor scale to obtain the co-modulation maps. To assess the statistical significance between conditions, we employed paired two-tailed t-tests with α =0.05

## RESULTS

### Ibogaine alters iEEG oscillatory components

To understand the acute effects of ibogaine on the rat brain, we recorded iEEG signals following its intraperitoneal administration (40 mg/Kg). Electrodes were located above the olfactory bulb (OB), primary motor (M1), primary somatosensory (S1) and secondary visual cortex (V2), allowing us to monitor the dynamical and regional effects of ibogaine (Figure 1A). As a working example, Figure 1B shows the OB time-frequency response after we administered saline (control) and ibogaine; time zero corresponds to the moment of injection. Compared to control, gamma oscillations (30-80 Hz) increased following the administration of ibogaine; this increase lasted for at least 2 hours. Note that this higher gamma power occurred associated with a longer time the animal spent awake (shown in the hypnogram). To analyze ibogaine effects at the group level, we considered only the wakefulness episodes in experimental and control conditions (Figure 1C). In comparison to control, ibogaine significantly increased gamma oscillations in the OB, M1, S1 and V2 areas (Figure 1C, and summarized in Figure 1D).

Along with the changes in gamma frequencies, the mean theta power increased (Figure 1D), while also decreasing its peak frequency from 9 Hz to 8 Hz (readily observed in S1 and V2 cortices because of their proximity to the hippocampus, Figure S1). Additionally, the high-frequency power (>100 Hz and up to 512 Hz) decreased in M1, S1 and V2 (see Figure 1C, D), though the lack of a spectral peak suggests this result arises from changes in muscular activity produced by the drug^27^.

### Ibogaine decreases inter-regional synchronization

Since ibogaine significantly altered the oscillatory power content of the iEEG, we next quantified its impact on long-range synchronization of brain areas within and across hemispheres (Figure 2A). Figure 2B shows an example of inter-hemispheric coherence between M1 cortices as a function of time (same animal as in Figure 1B). Interestingly, as opposed to its effect on gamma power, ibogaine strongly decreased inter-regional gamma synchronization.

Figure 2C shows a group level analysis separated by frequency bands by means of pair-wise electrode matrices (left column), which depict coherence differences (t-statistic) between conditions (saline vs ibogaine) in pseudocolor scale for each electrode pair (blue indicates a coherence decrease while red an increase). The electrode pairs with significant differences are also indicated in the right column. Ibogaine decreased phase coherence at the sigma-beta, gamma, and high-frequency bands in multiple cortical areas, including the OB, M1 and S1 (Figure 2C, D). In particular, inter-regional gamma coherence decreased in 9 of the 21 electrode pairs, including between right OB and right S1 cortex (Figure 2D, left panel), two areas that had an increase in their gamma power (Figure 1C). The same gamma coherence reduction occurred in the inter-hemispherical M1-M1 and M1-S1 electrode combination, but not in the intra-hemispherical M1-S1 (see Figure 2D and Figure S3).

### Ibogaine decreases iEEG temporal complexity

In the previous sections, we showed that ibogaine promoted local gamma oscillations which were uncoupled between areas. This activity resembles gamma oscillations that naturally occur during REM sleep^20,28,29^ (Figure S4), suggesting that the awake state under ibogaine exhibits similar REM sleep characteristics. To delve further into this matter, we tested the resemblance between states in their temporal complexity. This is important because the temporal complexity during REM sleep is significantly lower than during wakefulness, which can be observed independent of the cortical area and for a wide range of time-scales^30^.

In order to assess the temporal complexity, we down-sampled the original signals to 128 Hz (avoiding muscular contamination) and measured the permutation entropy of the time-series. This metric quantifies the diversity of dynamical motifs in the iEEG (larger values mean the signal has higher diversity, hence more complexity) and is robust to the presence of noise and short time measurements (see Material and Methods and ^24,31^). Figure 3 shows the average permutation entropy for each cortical electrode. Interestingly, in comparison to normal (control) wakefulness, ibogaine wakefulness displayed significantly lower levels of dynamical complexity in OB, M1 and S1 cortex. Note that these areas are the ones with most prominent changes in power and coherence. No significant changes were observed in V2. We should also point out that by virtue of downsampling, the gamma band oscillations are the only relevant frequencies contained in our complexity estimate.

**Figure 3.**
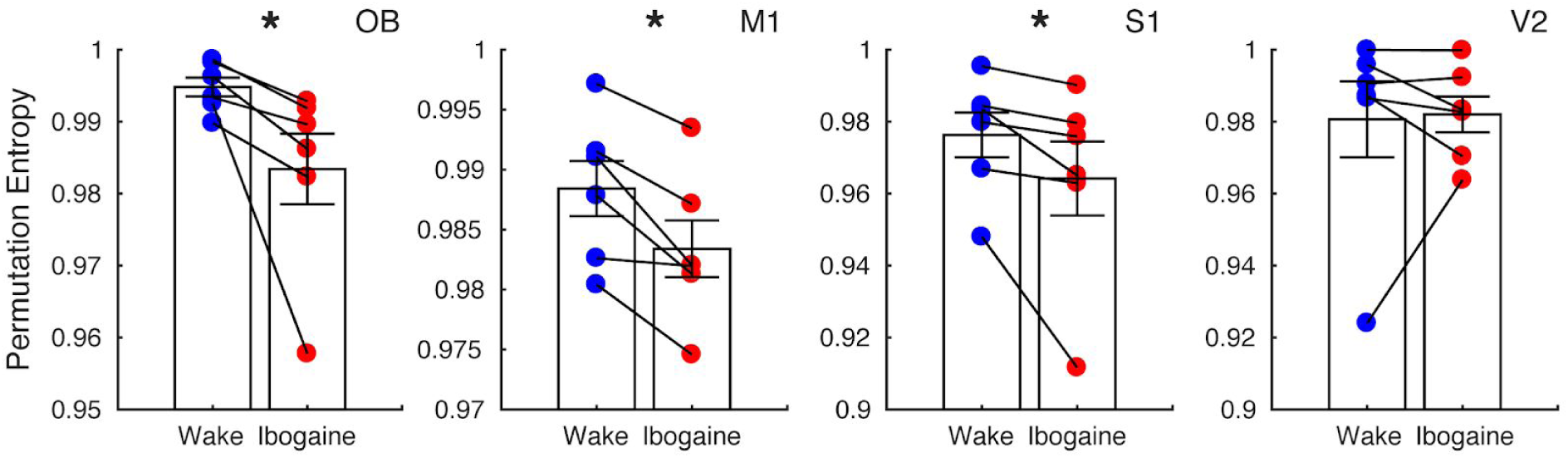
Ibogaine decreases iEEG complexity. Permutation entropy is employed to quantify the iEEG temporal complexity in normal (blue) and ibogaine (red) wake states (same electrodes as in Figure 1). Each dot shows the average permutation entropy of an animal (n = 6). Bars represent mean ± S.E.M.. *p<0.05, paired t-test.

### Ibogaine wakefulness and REM sleep have similar iEEG gamma activity

The previous section showed that ibogaine awake state differs from normal wakefulness. We next compared the ibogaine-induced brain state with physiological REM sleep (Figure 4A). We found that theta, sigma and beta power were lower during ibogaine wakefulness than in REM sleep. On the other hand, the high-frequency component (>100 Hz) had significantly higher power, likely due to muscular activity. Noteworthy, the power of gamma oscillations was similar between both states and minor statistically significant differences were found in the OB and M1 with larger gamma power during REM (Figure 4A). Furthermore, we also found similar levels of gamma coherence in the ibogaine wakefulness and REM sleep, even for electrode combinations which showed significant changes between physiological and ibogaine wakefulness (compare Figure 2B with Figure 4C). In contrast, the high-frequency spectrum was more coherent during ibogaine wakefulness than during REM sleep, probably as a consequence of the absence of muscle activity during REM sleep.

**Figure 4.**
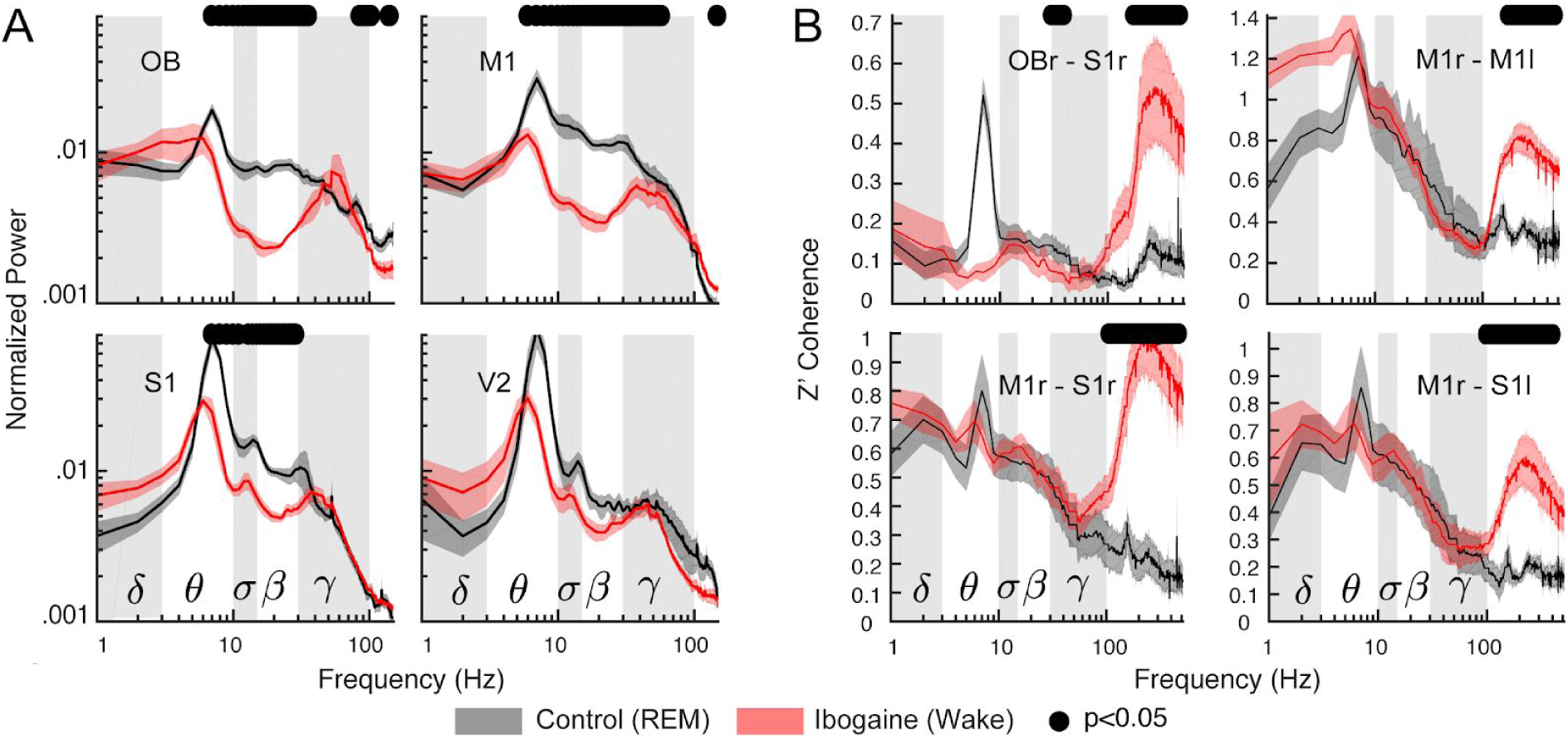
The ibogaine wakefulness shows REM sleep features in the gamma band. **A** Power spectrum comparisons between REM sleep (black) and the ibogaine wakefulness (red); only the right hemisphere is shown. The solid line represents the mean (n=6 animals); the shaded area depicts the S.E.M.. **B** Coherence comparisons between REM sleep and ibogaine. The black dots mark the statistically significant frequencies (p<0.05) corrected by a cluster-based permutation test

We also found that the temporal complexity during the ibogaine wakefulness was similar than during REM sleep (Figure S5); it was only significantly larger during ibogaine wakefulness in the M1 cortex. Overall, the data show that iEEG complexity values during ibogaine wakefulness are between normal wakefulness and REM sleep. Thus, although there are differences between ibogaine wakefulness and REM sleep, the power, complexity and inter-regional synchronization of gamma oscillations are comparable.

Finally, we directly tested whether the ibogaine wakefulness was closer to a REM-like state or to physiological wakefulness. For this purpose, we trained an artificial neural network to automatically classify the states of wakefulness and REM sleep. Figure 5A shows a schematic representation of the network, which is fed with the levels of gamma power and coherence of single 10-s artifact free epochs (input layer) and the output were the behavioral states (Wake or REM, output layer). After supervised training, the network successfully distinguished between wakefulness and REM (the confusion matrix for a representative animal is shown in Figure 5B). Then, the network was fed with ibogaine wakefulness data, these epochs were mostly classified as being REM sleep instead of wakefulness (Figure 5C). In fact, in 5 out of 6 animals the majority of the ibogaine epochs were classified as REM sleep, and in 3 animals all ibogaine epochs were classified as such. Therefore, these results show that the gamma oscillations induced by ibogaine have convincing REM sleep-like features.

**Figure 5.**
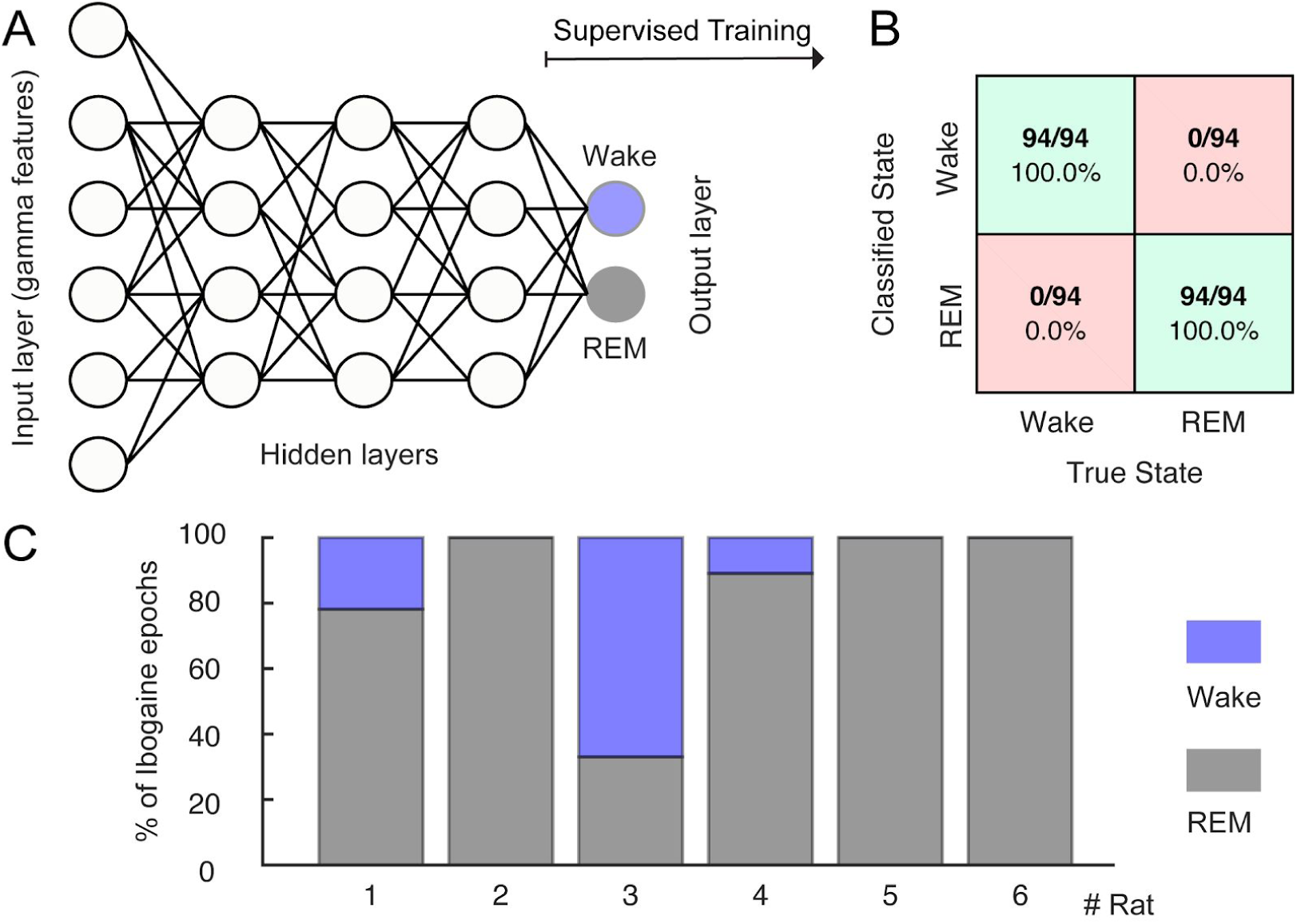
The ibogaine-induced brain state is considered a REM-like state by an automatic sleep scoring algorithm. **A** Schematic representation of the neural network employed to classify the states of wakefulness and REM sleep. The network contains one input layer which receives the gamma features (power and coherence of the OB, M1 and S1 electrodes), and 10 hidden layers (only 3 of them are shown in the picture). The network has one output layer with 2 nodes (Wake and REM). **B** Confusion matrix for an individual animal. **C** Bar plots showing how ibogaine wakefulness epochs were classified. Each animal is shown separately and was tested with its own control network.

## DISCUSSION

In the present study, we found that intraperitoneal administration of ibogaine in rats induces a waking brain state that has electrocortical REM sleep traits. These traits appear in the form of high power local gamma oscillations in the OB, M1, S1 areas, which are less coherent and less complex than in normal wakefulness. These features of gamma oscillations are similar to the ones present during REM sleep (Figures 4,5 and S4). Therefore, by measuring an important neurophysiological trait, our results support previous oneirogenic conjectures of ibogaine’s induced psychedelic state^6,16^. Interestingly, some of these traits were dragged into NREM sleep; compared to physiological NREM sleep, ibogaine NREM sleep showed gamma power increase circumscribed to the OB, and lower gamma coherence in several derivations (Figure S6).

When comparing our results to the effects elicited by other psychedelics, the lack of previous reports involving quantitative iEEG analysis of the psychedelic state in rodents, forces us to compare our results to previous literature in human beings. For instance, the administration of 5-HT_2A_ agonist (e.g. LSD, psilocybin) in humans reduces alpha (8-12 Hz) and beta band power and decreases their functional connectivity^32–35^. Similarly, our results show that ibogaine also reduced the connectivity at sigma and beta bands (10-30 Hz). Additionally, it is worth noting that we found significant changes in the OB, while in humans the predominant effects of traditional psychedelics are observed in the visual cortex^32^. Thus, both psychedelic effects involve major sensory areas relevant to each species. Furthermore, complementary analyses show that the gamma coupling to other frequencies is not affected by ibogaine in any of the cortical locations (Figure S7). Thus, as the slow OB oscillations (1-4 Hz) reflect the slow respiratory potentials^36,37^, our results suggest that sensory information is still likely to reach the OB, but is later integrated in an altered way, similar to the psychedelic state in humans^33^.

In addition to the electrophysiological similarities between ibogaine and serotoninergic psychedelics, the type of cognition elicited by the latter has been described as analogous to the one present during dreams^33^ (both referred as primary states of consciousness). In fact, a recent work shows that unlike other drugs (cocaine, opioids, etc), the semantic content of psychedelic experiences is closely related to dreams^38^. Thus, as dreams are to a large extent the cognitive correlates of REM sleep^39^, our report confirms such connection for ibogaine, and proves it by showing clear electrophysiological similarities between REM sleep and the psychedelic state induced by this drug.

Nevertheless, as mentioned before, human subjective reports also indicate differences between the experience elicited by ibogaine and classic psychedelics. Pharmacological and behavioural data in rodents also support these differences. While classical psychedelics share the ability to interact with the 5-HT_2A_ receptor in the low nanomolar range inducing the head twitch response (HTR) in rodents^11^, ibogaine binds to this receptor in the micromolar range (K _i_ 4.8 to 92.5 μM depending the study^40–43^) without producing HTR or similar responses^18^. Also, previous drug discrimination studies in rats showed that although ibogaine may produce some of its effect via 5-HT_2A_ activation (LSD and 2,5-Dimethoxy-4-methylamphetamine or DOM produced intermediate levels of substitution for ibogaine), this does not appear to be essential to the ibogaine-discriminative stimulus, since pirenperone (5-HT_2A_ antagonist) did not affect the ibogaine-appropriate response^43,44^. Further studies employing the same iEEG methodology used in this study will shed light into the electrophysiological similarities and differences between the wakefulness state induced by classical psychedelics and ibogaine.

When considering pharmacological interactions to explain the results obtained in this study, it should be noted that after i.p. administration, ibogaine is rapidly metabolized (half-life: 1.22 hs) to produce noribogaine, which has its own pharmacological and pharmacokinetic profiles^45,46^. According to our recently reported pharmacokinetic data^19^, both substances are present in the rat brain at pharmacologically relevant concentrations during the first two hours after ibogaine 40 mg/Kg i.p. administration, which corresponds to the time period studied in the present report. In this manner, ibogaine and noribogaine should be considered to explain our results. Although affinities displayed to most of the CNS receptors by ibogaine and noribogaine are low (micromolar range), we estimated high free drug maximum concentrations in rat brain for both substances in brain (∼ 3.1-3.5 μM)^19^. In this regard, noncompetitive antagonism at N-methyl-D-aspartate receptors (NMDA-R) by ibogaine (Ki ∼1.01-5.20 μM^40,47–53^) and in a less extent by noribogaine (Ki ∼ 5.48-31.4 μM^47,49,53^) should be considered as a key factor to explain the effects on the gamma band found on this study, since ketamine (a non-competitive NMDA-R antagonist) administration also produces a marked increase in gamma power^54–57^ while decreasing inter-regional gamma coherence^55,57^.

Nevertheless, effects on other neurotransmitter systems and receptors should be also considered. Since ibogaine and noribogaine inhibit serotonin re-uptake by modulating SERT activity (noribogaine being approximately ten-times more potent than ibogaine, IC50 for noribogaine ∼50−300 nM)^58,59^; the increase in the serotoninergic transmission, in addition to the above-mentioned interaction of ibogaine with 5HT_2A_ receptor, could explain some of the similarities found in the electrocortical activity between ibogaine and classic psychedelics. Further experiments are required to address the exact contribution of ibogaine and noribogaine (and their effects on neurotransmitter systems and/or receptors) to explain the results found on this study

As a final conceptual remark, REM sleep is considered a natural model of psychosis^60^ and in the late 1950’s psychedelics were studied as pharmacological models of psychosis^11^. Hence, considering the evidence provided in this study linking REM sleep to the brain-state induced by a psychedelic drug, it can be argued that REM sleep, psychosis and psychedelia are qualitatively and quantitatively part of a similar brain state. Which, in turn, can be achieved through different routes and measured from the electrocortical activity.

## Acknowledgments

This study was supported by the Programa de Desarrollo de Ciencias Básicas, PEDECIBA; Agencia Nacional de Investigación e Innovación (ANII), (FCE-1-2017-1-136550) and the Comisión Sectorial de Investigación Científica (CSIC) I+D-2016-589 grant from Uruguay. J.G was supported by CAP (Comisión Académica de Posgrado). N.R. acknowledges the CSIC group grant CSIC2018 - FID 13 - Grupo ID 722. A.B.L.T. was supported by Conselho Nacional de Desenvolvimento Científico e Tecnológico (CNPq) and Coordenacao de Aperfeicoamento de Pessoal de Nível Superior (CAPES), Brazil.

## Author contributions

J.G., M.C., I.C and P.T. designed the experiments; J.G., M.C. and A.M conducted the experiments; J.G., N,R. wrote analysis software; J.G. analyzed the data; J.G., M.C., A.M., S.C., N.R., A.B.L.T., I.C and P.T. were involved in the discussion and interpretation of the results; J.G., A.B.L.T., I.C and P.T. wrote the manuscript. All the authors participated in the critical revision of the manuscript, added important intellectual content, and approved the definitive version.

## Data and code availability

Data are available under request to the authors. The codes to obtain power and coherence spectra can be found in the Chronux toolbox and standard Matlab toolboxes. The code to perform the correction for multiple comparisons based on cluster-based permutation tests and to compute permutation entropy is freely available at: https://github.com/joaqgonzar/Ibogaine_analysis_2020

## SUPPLEMENTARY MATERIAL (1 Table and 6 Figures + text)

**Table S1.** Electrode Location.

| Electrode Label | Coordinates* |  |
| --- | --- | --- |
|  | Lateral | Antero-posterior |
| OBr | +1.25 mm | +7.5 mm |
| M1r | +2.5 mm | +2.5 mm |
| M1l | -2.5 mm | +2.5 mm |
| S1r | +2.5 mm | -2.5 mm |
| S1l | -2.5 mm | -2.5 mm |
| V2r | +2.5 mm | -7.5 mm |
| V2l | -2.5 mm | -7.5 mm |

\*All coordinates referenced to Bregma (Lateral: 0 mm, Antero-posterior: 0 mm)<sup>22</sup>.

**Figure S1.**
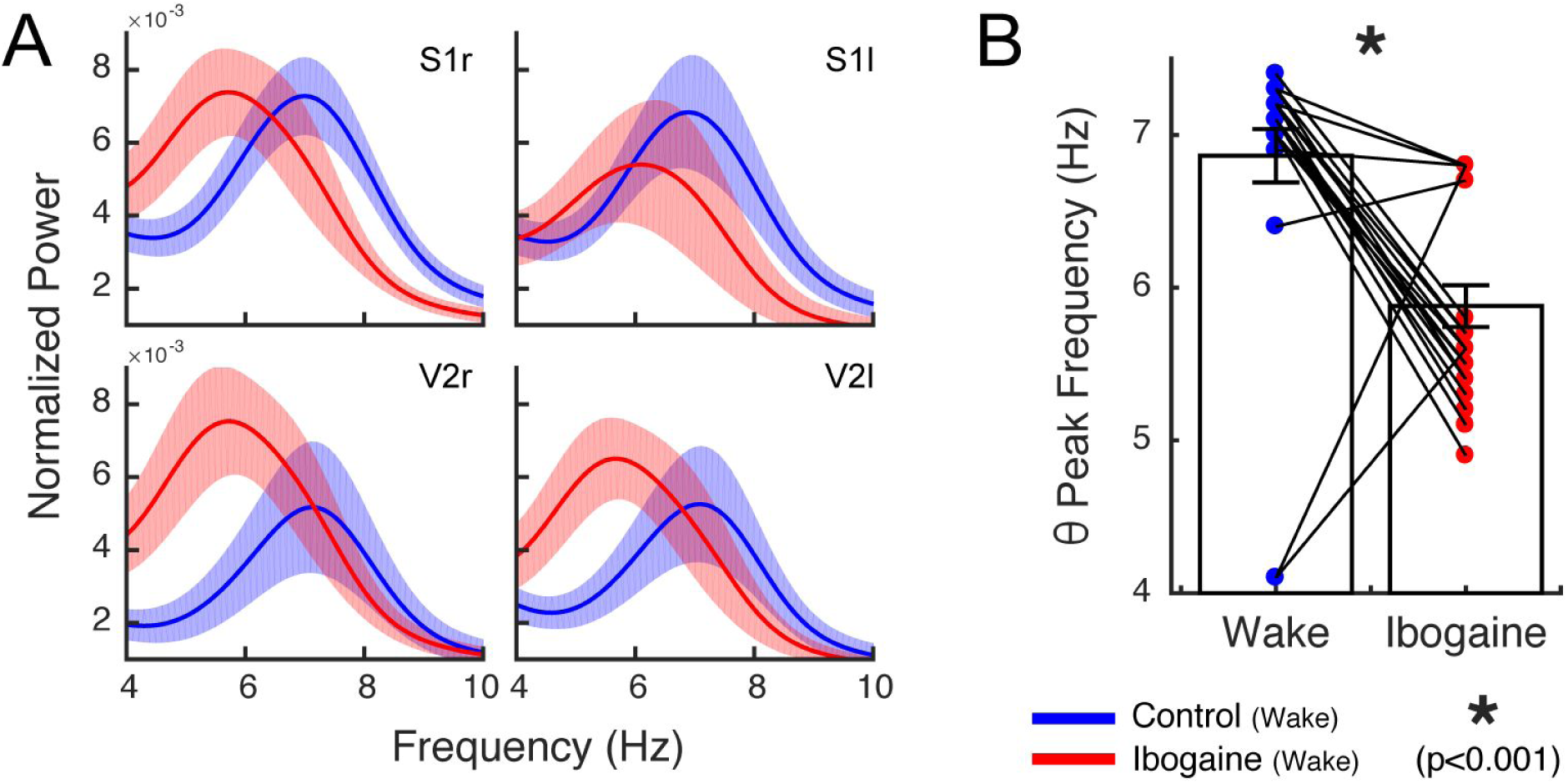
Ibogaine slows down theta peak frequency. **A** Normalized power spectrum in the theta range (4-10 Hz) for the four posterior electrodes. The solid line represents the mean (n=6 animals) for the first 2 hours post-injection; the shaded area depicts S.E.M.. Blue: control wakefulness. Red: ibogaine wakefulness. **B** Theta peak frequency during control and ibogaine wakefulness. Each dot is a cortical location of an individual animal (4 locations per animal). Bars represent mean ± S.E.M. *p<0.001, paired t-test.

**Figure S2.**
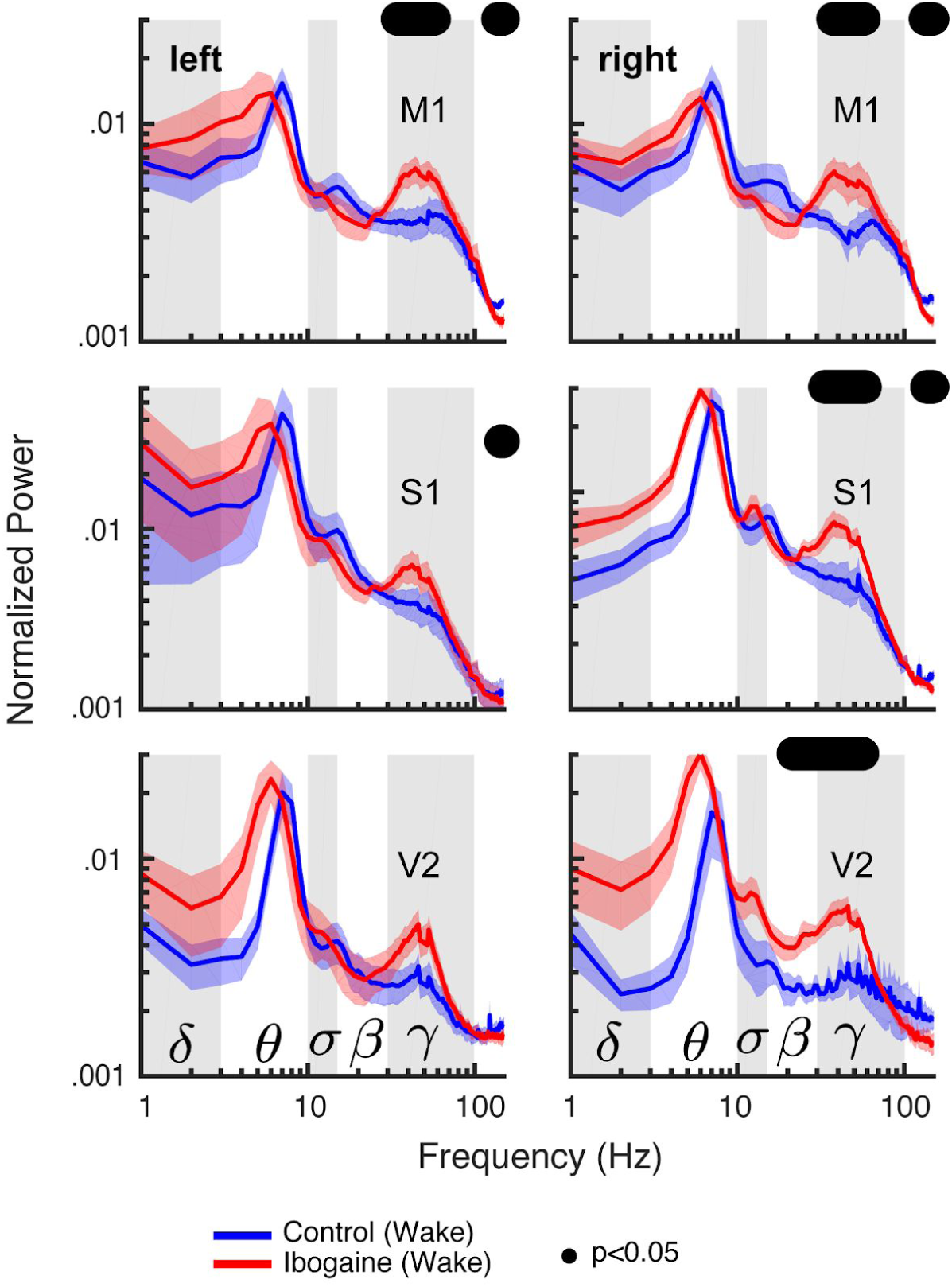
Ibogaine induces similar power changes in the left and right hemispheres. Blue: control wakefulness. Red: ibogaine wakefulness. The solid line represents the mean (n=6 animals) for the first 2 hours post-injection; the shaded area depicts S.E.M.. The black dots depict the statistically significant frequencies (p<0.05) corrected by a cluster-based permutation test. The spectra were whitened by multiplying power at each frequency by the frequency itself, and then normalized by dividing each frequency by the sum across frequencies.

**Figure S3.**
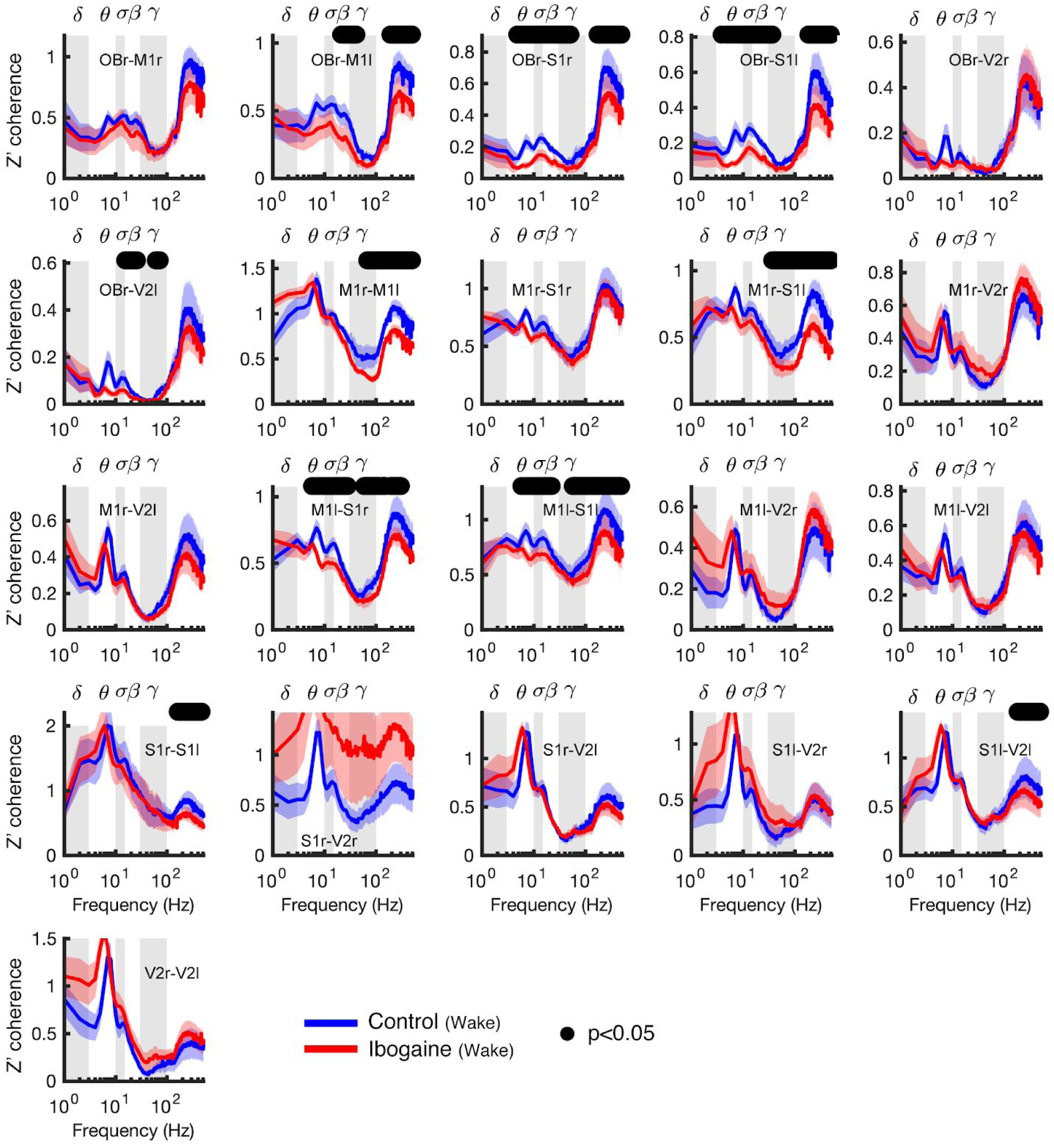
Coherence between all the electrode pairs during wakefulness. The solid line represents the mean (n=6 animals) for the first 2 hours post-injection; the shaded area depicts S.E.M. (r: right; l: left). The black dots depict the statistically significant frequencies (p<0.05) corrected by a cluster-based permutation test.

**Figure S4.**
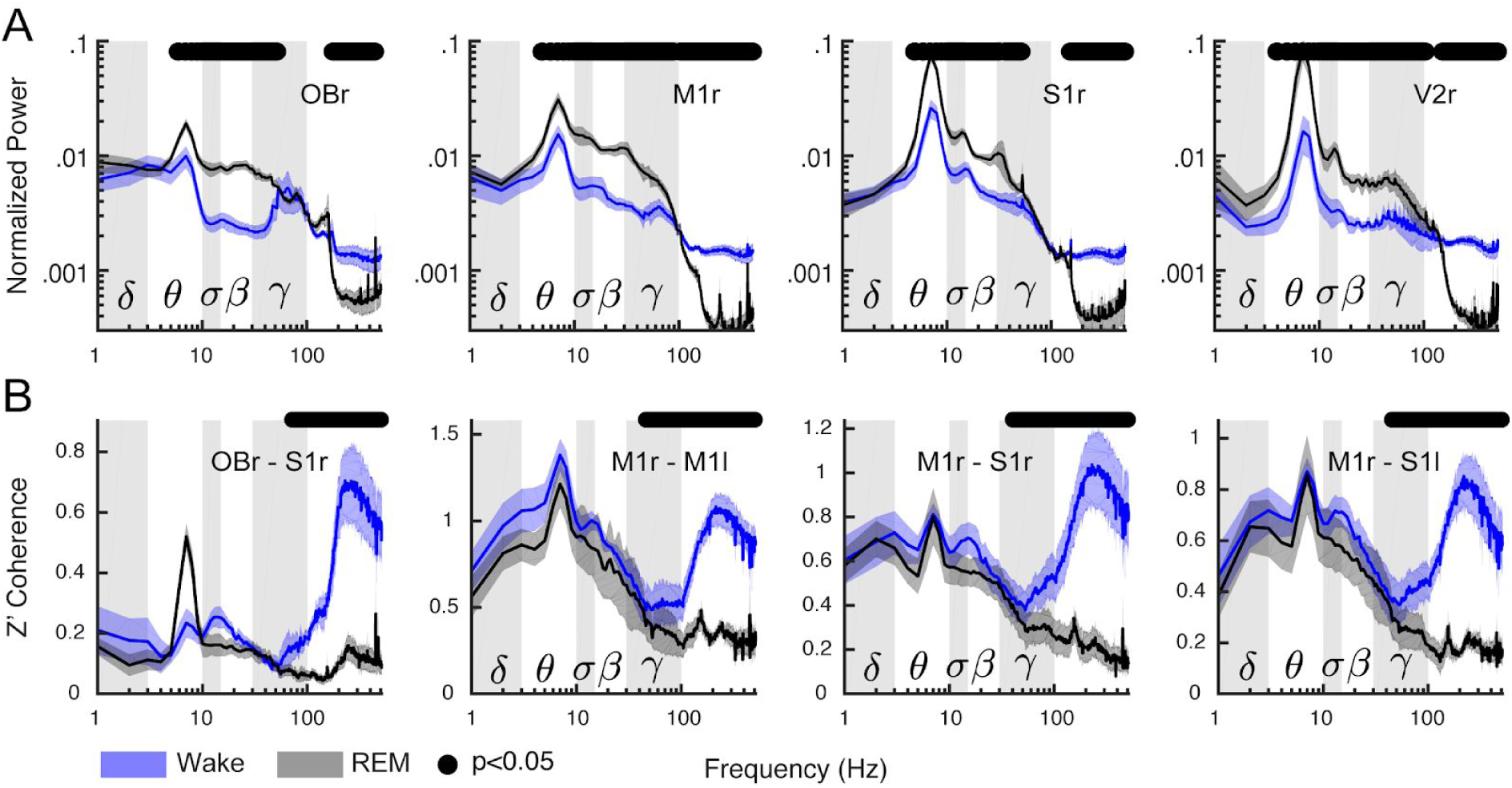
Gamma power and coherence during physiological wakefulness and REM sleep. **(A)** Power spectrum comparisons between physiological wakefulness (blue) and REM sleep (black); only the right hemisphere is shown. **(B)** Coherence spectra are shown for both normal wakefulness (blue) and REM sleep (black). The solid line represents the mean (n=6 animals); the shaded area depicts the S.E.M.. The black dots mark the statistically significant frequencies (p<0.05) corrected by a cluster-based permutation test (n=6 animals).

**Figure S5.**
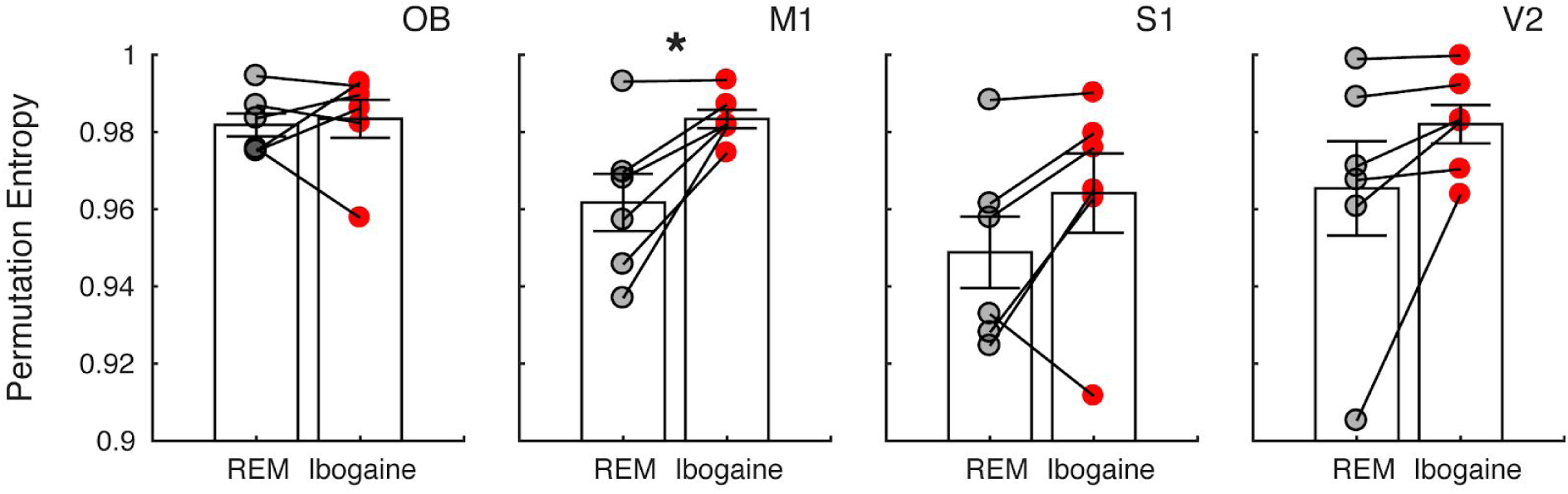
Permutation entropy during REM sleep and ibogaine wakefulness. Permutation entropy was employed to quantify the iEEG temporal complexity in physiological REM sleep (gray) and ibogaine wakefulness (red). Each dot shows the average permutation entropy of an animal (same electrodes as in Figure 1). Bars represent mean ± S.E.M.. *p<0.05, paired t-test (n=6 animals).

**Figure S6.**
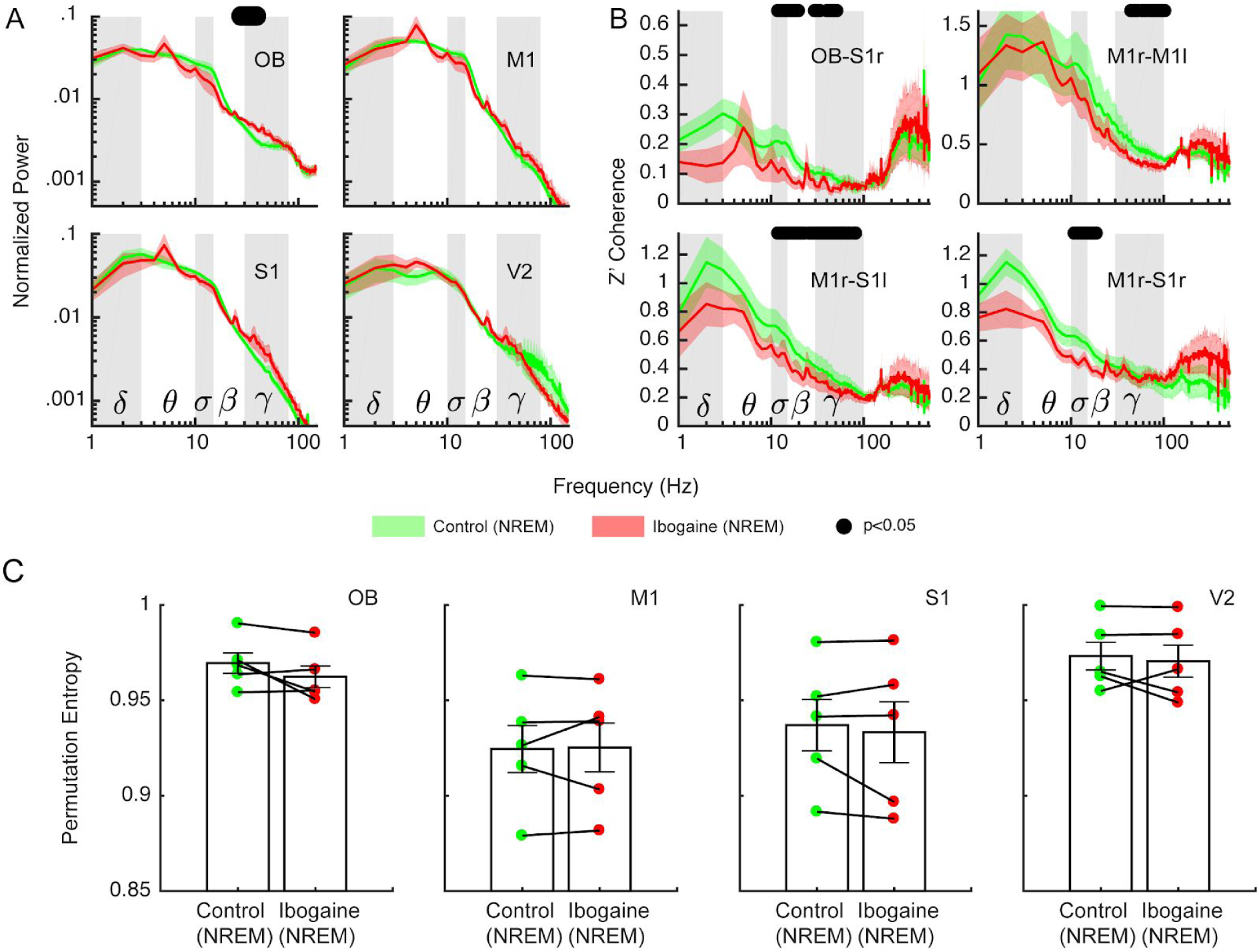
Ibogaine decreases inter-regional synchronization during NREM sleep. **A**,**B** Normalized power (**A**) and coherence (**B**) spectra for the physiological (green) and ibogaine NREM sleep (red). The solid line represents the mean (n=6 animals) for all NREM epochs within the first 6 hours post-injection; the shaded area depicts S.E.M. (r: right; l: left). The black dots depict the statistically significant frequencies (p<0.05) corrected by a cluster-based permutation test. **C** Permutation entropy comparisons between physiological NREM sleep (Control) and ibogaine NREM sleep (Ibogaine). Each dot shows the average permutation entropy of an animal (same electrodes as in Figure 1). Bars represent mean ± S.E.M.. No significant differences were found, paired t-test (n = 5 animals, one animal did not reach the minimum time required for the analysis).

**Figure S7.**
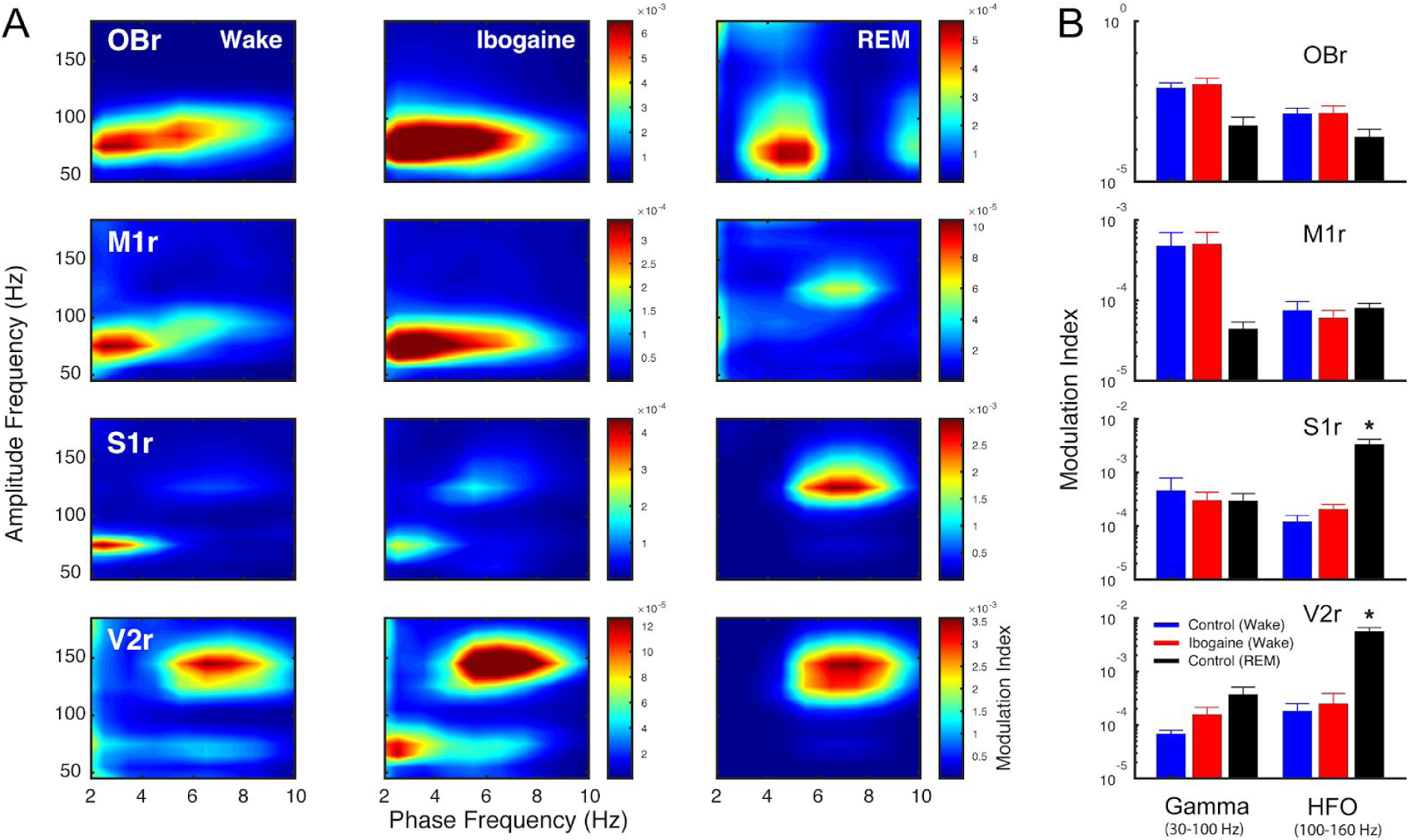
Gamma band cross-frequency coupling remains unaltered after ibogaine administration. **(A)** Population averaged co-modulation maps for control wakefulness, ibogaine wakefulness and REM sleep. Red colors indicate high phase-amplitude coupling (see Material and Methods). **(B)** Modulation index for theta-gamma (30-100 Hz) and theta-high-frequency oscillation (100-160 Hz, also known as the fast-gamma band) coupling. For each animal and state, the maximal modulation index value within the analyzed frequency ranges was taken for each cortex. Bars represent mean ± S.E.M. over animals. *p<0.05 paired t-test against ibogaine wakefulness state (n = 5 animals, one animal had large co-modulation artifacts).

